# Hippocampal granule cells balance energy and function along the Pareto optimal front

**DOI:** 10.64898/2026.09.29.755037

**Authors:** Martin Mittag, Alexander D. Bird, Lucas A. Mongiat, Arnd Roth, Hermann Cuntz, Peter Jedlicka

**Affiliations:** ICAR3R – Interdisciplinary Centre for 3Rs in Animal Research, Justus-Liebig-University Giessen, 35390 Giessen, Germany; Institute for Anatomy II, Dr. Senckenberg Anatomy, Neuroscience Center, Goethe University Frankfurt, Frankfurt am Main, Germany; Medical Physics Department, Centro Atómico Bariloche - CNEA, Argentina; Wolfson Institute for Biomedical Research, University College London, London, UK; Ernst Strüngmann Institute (ESI) for Neuroscience of the Max Planck Society, 60528 Frankfurt am Main, Germany

**Keywords:** Compartmental modelling, Dentate gyrus, Pattern separation, Energy efficiency, Multi-objective optimisation, Adult neurogenesis

## Abstract

The dentate gyrus’ principal neuron, the granule cell (GC), plays a key role in pattern separation, transforming similar inputs into dissimilar outputs. We used a new measure based on information theory to evaluate this transformation in an experimentally derived *in silico* GC model with realistic ion channel distributions across its dendrites. We found that model performance aligned well with current-clamp recordings from mouse brain slices. However, it was also possible to find artificial parameter combinations in the model that functionally outperformed the recorded cells while matching their action potential shapes. Rather than maximising computational power, the biologically realistic GC model balanced energy efficiency against increased functionality in a Pareto optimal way: No possible parameter combination was found to improve one of the objectives without impairing the other. Interestingly, immature GCs adopted a different weighted trade-off with a stronger emphasis on energy efficiency than pattern separation, but they remained Pareto optimal over their putative developmental trajectory. Our results support the idea that evolution favours neurons that can satisfy functional and energetic demands simultaneously.

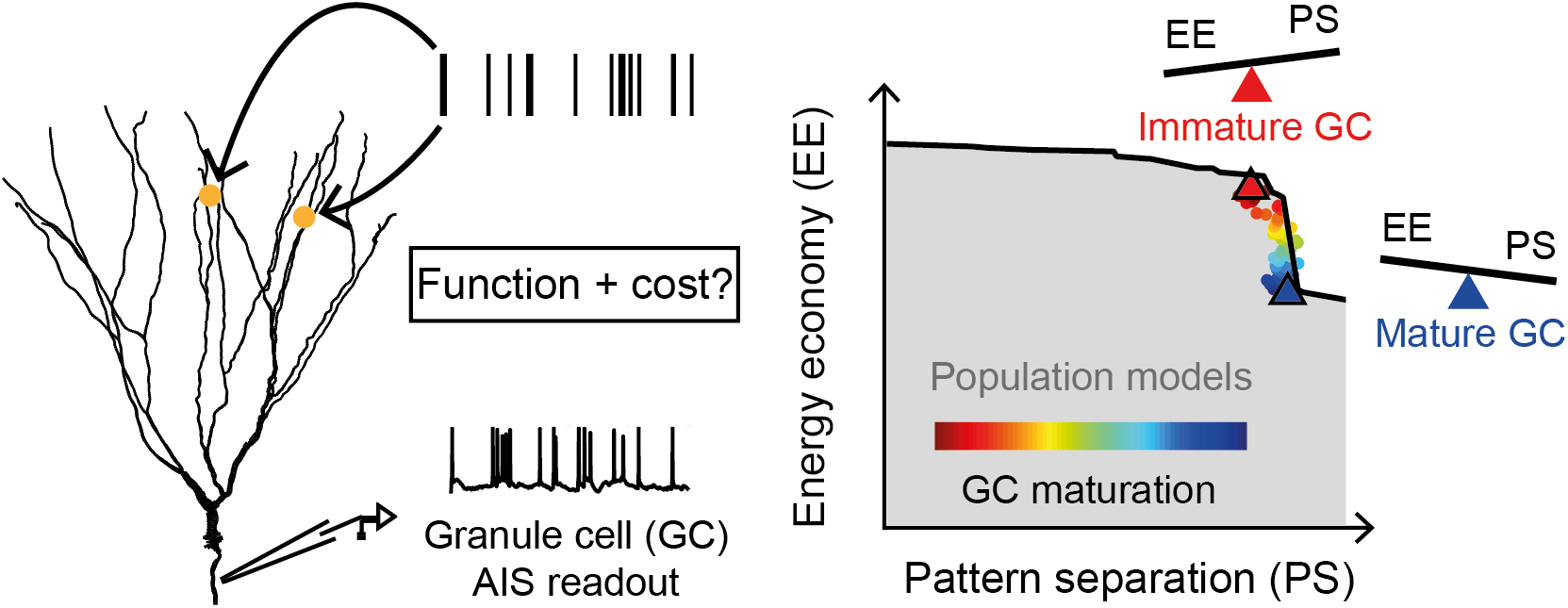

**In brief:** We find that dentate gyrus granule cells are Pareto optimal for the simultaneous performance of pattern separation and energy economy, adopting a differently weighted trade-off depending on their developmental stage.

**Highlights:**

- An experimentally constrained single granule cell (GC) model can perform temporal pattern separation in line with real data.
- Artificial channel configurations can improve pattern separation, but strictly at the cost of increased ATP consumption.
- GCs with realistic channel configurations appear Pareto optimal for the simultaneous performance of pattern separation and energy economy.
- Immature GCs remain on the Pareto front but prioritise low energy expenditure over pattern separation.

## Introduction

Optimisation is a powerful conceptual framework to understand the intricacies of biological function. By identifying what a given structure is optimised to do, one can make inferences about the evolutionary pressures on its real-world roles (Hart *et al*., 2015). This approach is especially potent for highly complex systems with multiple competing objectives, such as neurons (Pallasdies *et al*., 2021; Jedlicka et al., 2022; Laughlin and Sejnowski, 2003). Neurons feature numerous ion channel proteins distributed across branched dendrites to regulate input responses (Bast *et al*., 2023). The extent to which biophysical and structural optimisation explains single-neuron computation remains an open question.

To address this, we focus here on hippocampal granule cells (GCs), for which a particularly detailed computational multi-compartmental model exists that reproduces a large number of experimental and pharmacological manipulations (Beining *et al*., 2017b). A major candidate for GC function is pattern separation, the transformation of similar inputs to highly dissimilar outputs (Marr, 1971; Leutgeb et al., 2007; Bakker et al., 2008; Yassa and Stark, 2011; Neunuebel and Knierim, 2014; Chavlis and Poirazi, 2017). In this way, GCs contribute to the learning and memory capacity of the hippocampus (Marr, 1969; Albus, 1971). Though primarily a network phenomenon (Cayco-Gajic and Silver, 2019), individual GCs participate via significant temporal filtering (Madar *et al*., 2019; Lö ffler and Gupta, 2022). Crucially, computation incurs a high energetic cost. Because brain energy supply heavily limits neuronal activity (Attwell and Laughlin, 2001; Laughlin and Sejnowski, 2003; Padamsey et al., 2022), improved computational performance in the form of enhanced information processing often demands higher metabolic expense (Harris *et al*., 2015; Remme et al., 2018; Sommer et al., 2026). This imposes strong constraints on the optimality of neurons for computational function.

Two biological phenomena impact the direction of GC function trade-offs and thus make GC function the ideal test subject for a study of optimal computation: ion channel degeneracy and adult neurogenesis. GC models exhibit degeneracy, where distinct channel combinations yield similar somatic electrophysiology (Schneider *et al*., 2023). Additionally, the dentate gyrus continuously generates new GCs (Altman and Das, 1965; Kuhn et al., 1996) that integrate into the circuit as they mature (Ge *et al*., 2007; Beining et al., 2017a; Groisman et al., 2020). Immature GCs are highly excitable (Mongiat *et al*., 2009; Mongiat and Schinder, 2011) with weak, non-specific inputs (Toni *et al*., 2007; Vivar et al., 2012; Restivo et al., 2015). Their computational role, whether aiding pattern separation (Sahay A *et al*., 2011; Nakashiba et al., 2012a), pattern completion (Kempermann, 2002; Finnegan and Becker, 2015), or learning new patterns (Aimone *et al*., 2011; Johnston et al., 2016; Danielson et al., 2016), remains to be fully understood during GC development.

Here, we investigate whether individual GCs optimally perform pattern separation under energy cost constraints and how this optimisation is supported by ion channel degeneracy. We explore whether pattern separation and energetic economy are opposing or complementary objectives, and how they shape the parameter space of mature GCs. Finally, we study how the distinct biophysical properties of immature GCs alter the weighting of the two-objective computational performance.

## Results

### GCs are not optimal for pattern separation alone

In order to explore GC function, we applied our recently developed information-theoretic pattern separation measures (Bird *et al*., 2024) to both existing mouse GC patch-clamp data (Madar *et al*., 2019) and a state-of-the-art biophysically realistic and experimentally validated *in silico* GC model (short: bvGC model, Beining *et al*., 2017b). Perforant path inputs were stimulated and GC spiking responses were recorded across a range of input correlations (10*Hz*, five spike trains within the ensemble, 10 repetitions, correlations varied between 0.05 − 1; **Fig 1A**). Specifically, our previously validated measures (Bird *et al*., 2024) - relative redundancy reduction and sparsity-weighted mutual information (MI) - evaluate the transformation of similar input spike patterns into dissimilar output patterns while penalizing information loss. This ensures that pattern separation performance is not trivially enhanced by the deletion of informative spikes. For both measures, pattern separation increased with input correlation (**Fig 1B**, black). However, there was a decline at an input correlation of approximately 0.975, indicating that noise impairs pattern separation close to, but not at, fully correlated input patterns.

**Figure 1.**
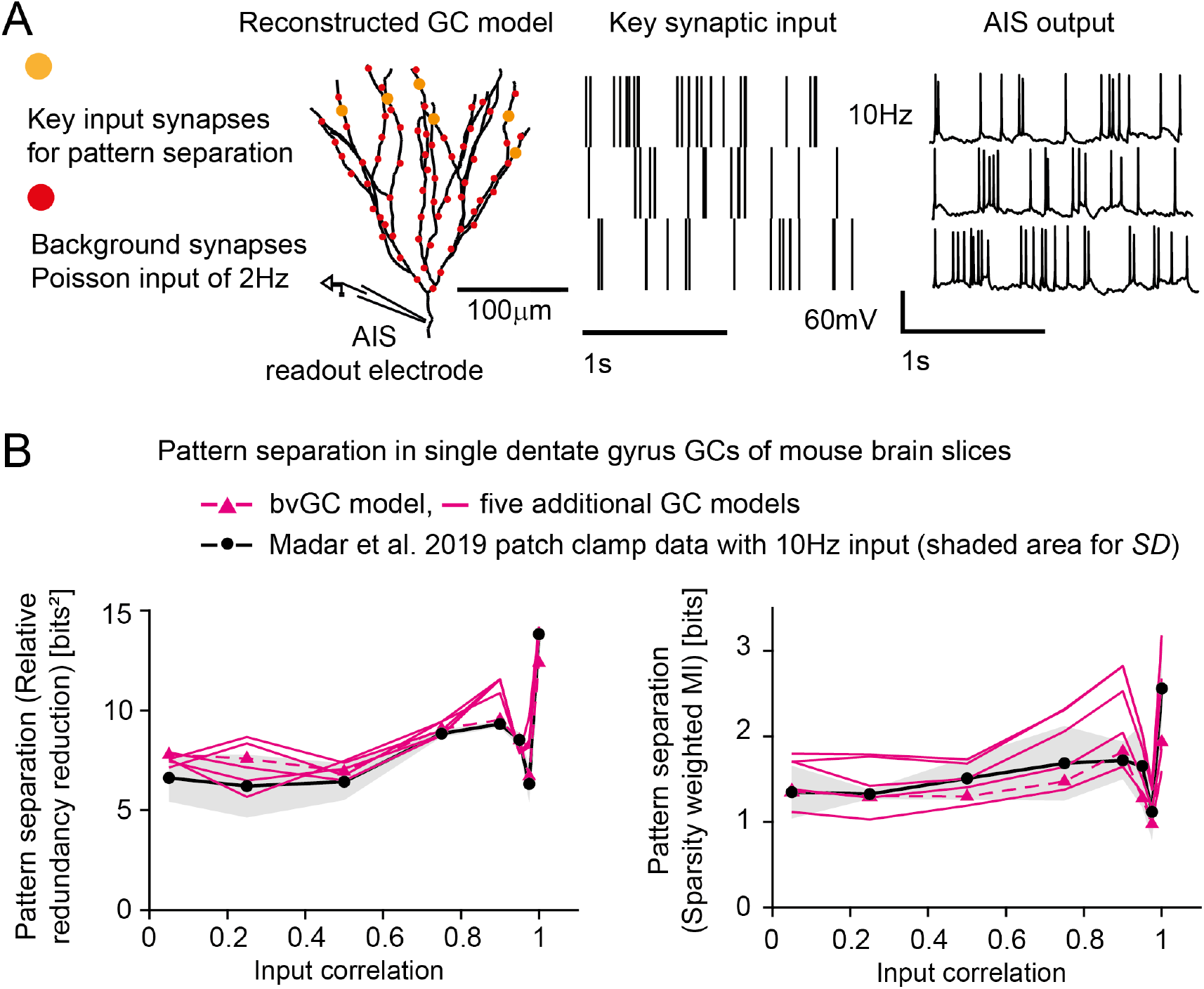
Comparison of a biologically validated GC model (bvGC) to experimental data for newly established pattern separation measures. **A**, *Left*: Example GC morphology shown without diameters for better visualisation and key input synapses (*orange*, compare background synaptic activity in *red*) for pattern separation, *Middle:* 10*Hz* input spike ensemble, *Right:* corresponding axon initial segment (AIS) output traces. **B**, *Left*: Pattern separation quantified via relative redundancy reduction (Bird *et al*., 2024) in single dentate gyrus GCs of mouse brain slices (*black dots*, Madar *et al*., 2019, shaded area for standard deviation) compared to various GC models (bvGC *dashed pink*, additional models *solid pink*), *Right*: same for pattern separation quantifed via sparsity weighted mutual information (Bird *et al*., 2024).

The bvGC model performance fell within the range of the experimental data (**Fig 1B**, dashed pink). Interestingly, for traditional pattern separation measures that do not penalise information loss, such as decorrelation or orthogonalisation, the performance of the experimental data deviated from the bvGC model (**Fig S1A**). This indicated that our information-theoretic pattern separation measures capture important aspects of real GC function.

Surprisingly, it was possible to generate a degenerate population of GC models with different biophysical parameter values, but the same somatic responses to current injection (Schneider *et al*., 2023), where some parameter combinations outperformed pattern separation from experimental GC measurements (**Fig 1B**, solid pink). This strongly indicated that real GCs had further objectives beyond only optimising the pattern separation computation.

### Economy of energy as a second objective

GCs are highly economical in terms of energy consumption. They are electronically compact, sparse-firing and possess extremely metabolically efficient action potential biophysics (Alle *et al*., 2009). Whilst it is possible to experimentally probe individual aspects of energy consumption, an established ion-counting method (Remme *et al*., 2018) in detailed biophysical models allows one to estimate the metabolic demands of an entire GC during realistic activity. Here, we counted the total inward currents of sodium (Na^+^) and calcium (Ca^2+^) and calculated the ATP cost to extrude these ions through transmembrane pumps (Attwell and Laughlin, 2001). We first sought to assess and validate the energetic properties of our bvGC and population models, which were originally neither tuned nor selected for economy, against the available experimental literature.

A major determinant of neuron action potential efficiency is the overlap of sodium (Na^+^) influx and potassium (K^+^) efflux (Crotty and Levy, 2007; Carter and Bean, 2009; Sengupta et *al*., 2010; Stemmler et al., 2011; Niven, 2016; **Fig 2A**). The mossy fibre boutons (MFBs) of GCs have been shown to have a charge overlap that is far closer to the theoretical minimum for action potential propagation than that found in classical Hodgkin-Huxley models based on squid giant axons (Hodgkin *et al*., 1952; Alle et al., 2009).

**Figure 2.**
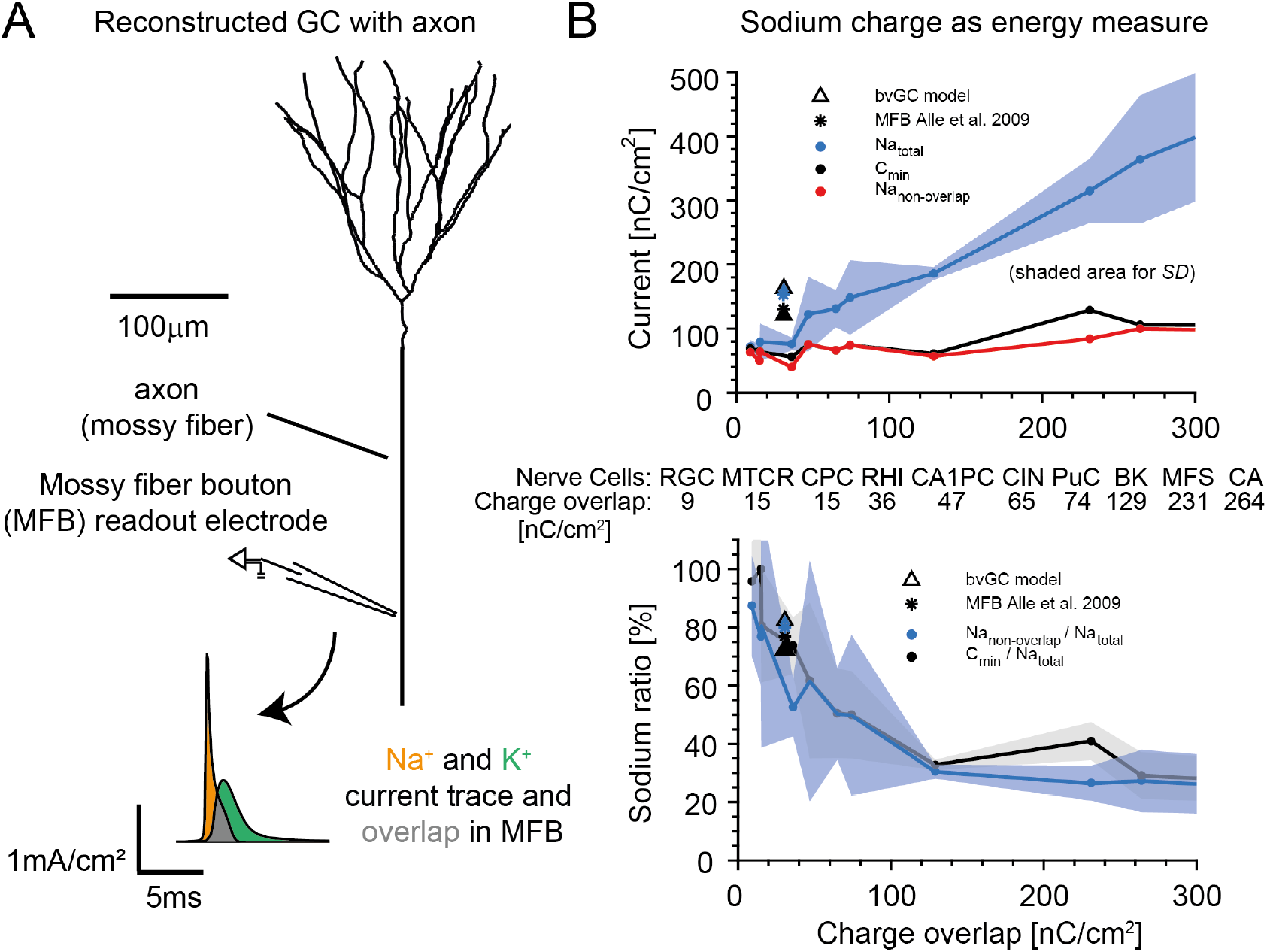
The bvGC model displays realistic sodium charge and ratio as energy measures in MFBs of GCs and other nerve cells. **A**, Reconstructed GC morphology with mossy fiber bouton (MFB). Diameters were left out for better visualisation. The inset shows the sodium (*orange*) and potassium (*green*) currents and their overlap (*grey*) recorded from the MFB. **B**, *Top*: Comparison of total sodium (*blue*, shaded area for standard deviation), capacitive minimum current (*black*) and non-overlapping sodium (*red*) with respect to the charge overlap of sodium and potassium in various nerve cells (with increasing charge overlap: rat granule cell RGC, mouse thalamo-cortical relay neuron MTCR, cortical pyramidal cell CPC with *n* = 8 cells, rat hippocampal interneuron RHI, CA1PC with *n* = 8 cells, cortical interneuron CIN with *n* = 5 cells, purkinje cell PuC with *n* = 7 cells, honeybee Kenyon cell BK, mouse fast-spiking neuron MFS, crab motor neuron axon CA; see also the inset table; for modelled cells no *n* is mentioned), including the bvGC model (*triangle*, Beining *et al*., 2017b) and MFB data (*star*, Alle *et al*., 2009). *Bottom*: Data as above. Ratio of non-overlapping sodium (free sodium with respect to potassium) and total sodium as energy efficiency measure.

To juxtapose the energetic properties of the bvGC model with a range of experimental findings, in **Figure 2B** and **Table S1** we summarized existing data on various nerve cells (CA1 PC with *n* = 8 cells, cortical interneuron CIN with *n* = 5 cells, purkinje cell PuC with *n* = 7 cells and cortical pyramidal cell CPC with *n* = 8 cells from Carter and Bean, 2009; and models of crab motor neuron axon CA, mouse fast-spiking neuron MFS, honeybee Kenyon cell BK, rat hippocampal interneuron RHI, rat granule cell RGC, mouse thalamo-cortical relay neuron MTCR as in Sengupta *et al*., 2010) including the MFB of the GC (Alle *et al*., 2009). We compared this data to the bvGC model (Beining *et al*., 2017b). **Figure 2B** top shows the expected linear increase of total Na^+^ charge with increasing charge overlap (*blue*). Free Na^+^ and the minimum capacitive current needed for the voltage change during the firing of the action potential remain relatively constant with increasing charge overlap (*red* and *black*). However, the ratio of free and total sodium charge falls exponentially with increasing charge overlap (**Fig 2B**, *bottom*). Crucially, the bvGC model showed strong agreement with the MFB data (*star*), for which it was not fitted, providing evidence that the model reflects the true metabolic costs of GC firing.

### The bvGC model lies on the Pareto front for energy economy and pattern separation

We have shown that pattern separation and energy economy are both important objectives for GCs. To investigate both at the same time, we applied the multi-objective (Pareto) optimality framework to the degenerate population of GC models, members of which have indistinguishable voltage responses to somatic current injection despite different underlying biophsyical parameters (c.f. Alon, 2006; Sheftel et al., 2013; Jedlicka et al., 2022; Schneider et al., 2023; **Fig 3A**). Phase-locked spike train ensembles with three component trains were fed to the GC outer molecular layer at individual firing rates of 10Hz, 15Hz and 20Hz and a 6Hz *theta*-band phase rate, representing lateral perforant path inputs (Quintanilla *et al*., 2022). Plotting ATP consumption against information transmission and pattern separation performance revealed that, despite the models being degenerate to somatic current inputs, they displayed very different responses to distributed synaptic inputs with respect to pattern separation and its energy cost (**Fig 3B**). In particular, models that outperformed the real GCs at pattern separation (**Fig 1B**) did so at a high energetic cost compared to the rest of the population (**Fig 3B**, *Magenta*).

**Figure 3.**
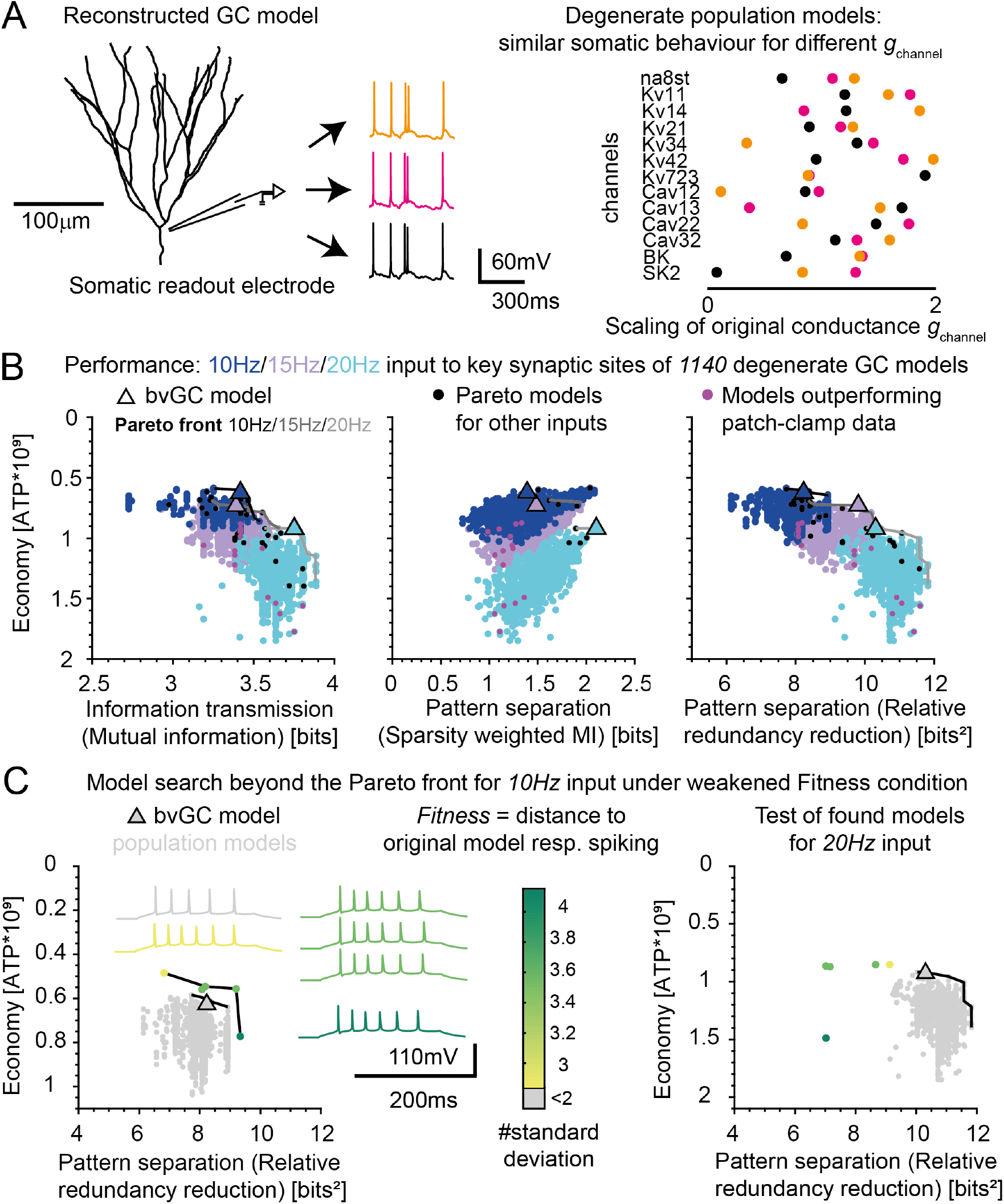
The bvGC model robustly maintains Pareto optimality for pattern separation and energy economy across a population of degenerate GC models. **A**: Example of a reconstructed GC morphology. Diameter neglected for better visualisation (*left*). Three similar somatic voltage traces (*middle*) corresponding to three (*orange, pink, black*) different sets of channel parameters (*right*). **B**: Simultaneous performance of economy of energy and pattern separation (*left* MI, *middle* sparsity weighted MI, *right* relative redundancy reduction) in a population of degenerate GC models for three different input conditions (*blue* 10*Hz, magenta* 15*Hz, cyan* 20*Hz*). The bvGC model (*triangle*) is close or on the Pareto front (*black* 10*Hz, grey* 15*Hz, light grey* 20*Hz*) in all conditions. Models that were Pareto optimal for the other input conditions are shown in *black dots* while models that outperformed data from Madar et al. (2019) are shown in *magenta dots*. **C**: *Left*: Models beyond Pareto front (*black line*) for relaxed fitness condition during 10*Hz* synaptic stimulation. The degenerate population models and the bvGC model (*triangle*) have a *fitness <* 2 (*grey*), while the found models beyond the Pareto front showed impaired fitness (*yellow, light green, green*); see also the example traces with corresponding colour in the inset. *Right*: Same models as left (*grey, yellow, light green, green*) but for 20*Hz* input.

A subset of the models was Pareto optimal in this population: they could not be improved in terms of energy economy without losing pattern separation performance and vice versa. Strikingly, the bvGC model (Beining *et al*., 2017b), which was not fitted for energy economy or pattern separation performance but constrained by experimentally measured ion channel densities, was on or close to the Pareto front for all input conditions (**Fig 3B**, *triangle*; see also **Fig S2** for a population of degenerate GC models with axons). Importantly, other models that were Pareto optimal for one input condition were typically not optimal for other input frequencies(**Fig 3B**, *Black*). This indicates that the bvGC model is more robust and maintains Pareto optimality for a wider range of inputs.

The bvGC model is Pareto optimal in the degenerate population, but the question remained as to whether it is possible to improve functional performance by allowing voltage responses to deviate from the data. **Fig 3C** shows that it is possible to obtain functionally better cells for a specific input (i.e. 10*Hz*) with spike shapes and responses that differed from those observed experimentally. However, these fitted, unrealistic cells perform very poorly for inputs different from those they were fitted to (**Fig 3C**, *right*, 20*Hz*). This implies that the biophysics of real GCs confer robustness to different input patterns in the trade-off between energy economy and pattern separation performance.

### Ion channels supporting Pareto optimal GC function

Having established that biophysically realistic GC models are Pareto optimal across the two tasks of pattern separation and energy economy, we sought to understand how their specific ion channel distributions support their performance. Economy (estimated as negative ATP count) correlated positively with the axon hillock *K*_*v*_7.2*/*3 channel, somatic *K*_*v*_2.1 channel and most strongly with the dendritic *K*_*v*_4.2 channel (**Figure 4A**). Conversely, a negative correlation was seen for the axon hillock *na*8*st, K*_*v*_1.4 and *K*_*v*_3.4 channels, indicating that channels involved in spike generation are energetically costly. In the 20Hz condition, pattern separation showed a positive correlation with the axon hillock *na*8*st* and *K*_*v*_1.4 channels in contrast with their negative relationship with energy economy.Additionally, the *K*_*v*_3.4 channel correlated negatively with both pattern separation and economy. Interestingly, in addition to its positive correlation with energy economy, the dendritic *K*_*v*_4.2 channel also exhibited a mild positive correlation with pattern separation, suggesting that this conductance synergistically supports both objectives.

**Figure 4.**
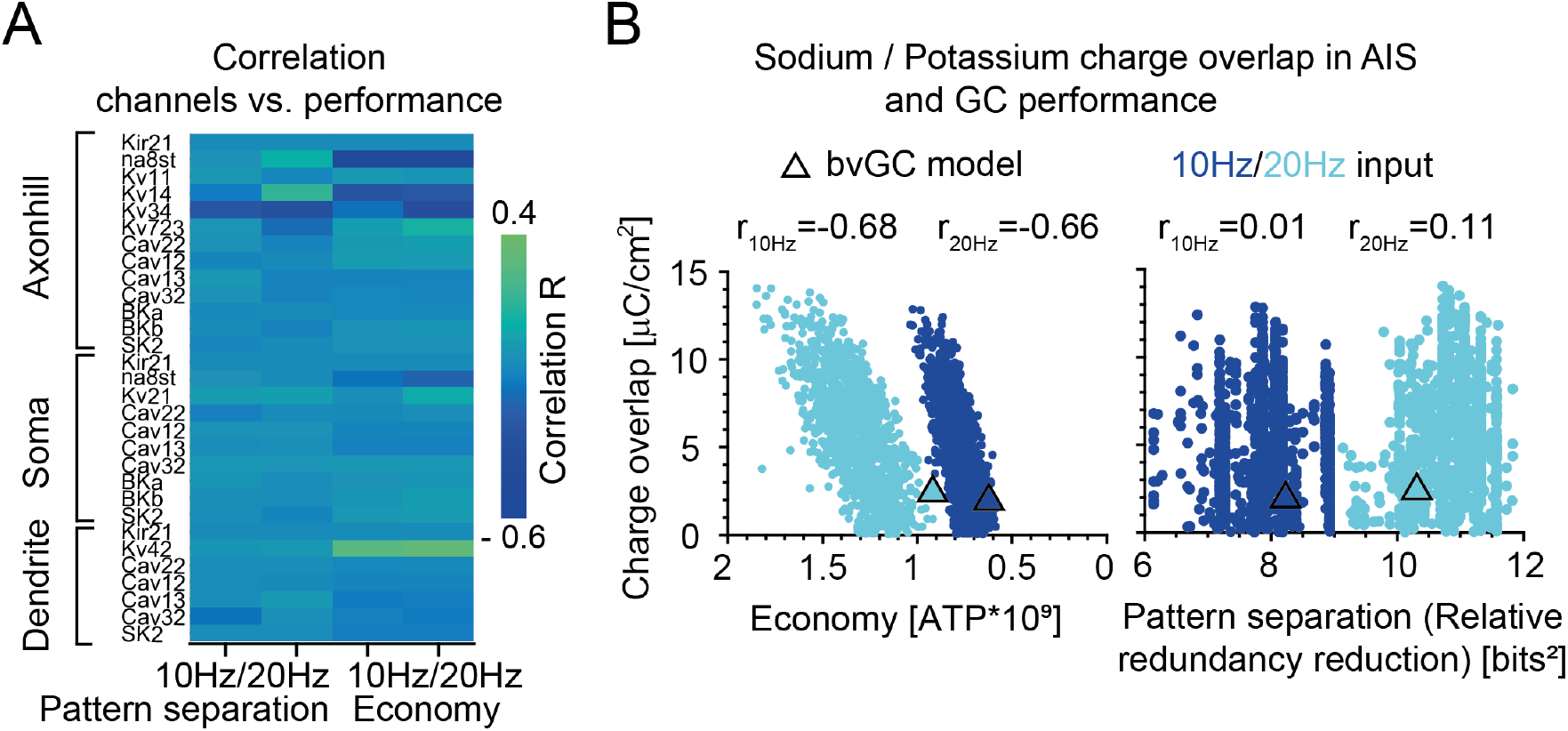
Ion channel mechanisms underlying the performance for pattern separation and its energy economy in a population of degenerate GC models. **A**: Correlation coefficient *R* (negative *blue*, positive *green*) between the channel conductances (30) versus GC economy and pattern separation in the population models. **B**: Sodium-potassium charge overlap in the AIS versus economy and pattern separation for two input conditions of 10*Hz* (*dark blue*) and 20*Hz* (*cyan*). The bvGC model is depicted with a *triangle*. The correlation values for both input frequencies (*r*_10*Hz*_, *r*_20*Hz*_) are indicated.

Regarding spike generation, we observed a negative correlation between sodium and potassium charge overlap during action potentials in the axon initial segment and total energy economy (*r*_10*Hz*_ = −0.68, *r*_20*Hz*_ = −0.66, **Fig 4**, *right*). Thus, as expected, more energy efficient action potentials imply more economical neurons. However, no correlation appeared between charge overlap and pattern separation, indicating that spike generation biophysics are largely independent of computational power for these inputs.

Taken together, although extrinsic (synaptic) properties also play a role (see below), these results show that the trade-off between pattern separation and energy economy in GC models is also linked to the distribution of intrinsic ion channels. While some channels, like *K*_*v*_4.2, synergistically push GC models toward optimal two-objective performance, others, such as *na*8*st* and *K*_*v*_1.4, show mostly opposing effects across the two objectives. We predict that channels that correlate with both objectives move the GC performance toward the Pareto front. In contrast, channels that display opposing correlations for pattern separation and economy influence where the GC resides along the Pareto front (see also below). Because GCs cannot improve pattern separation by changing these channel densities without compromising energy economy, realistic GC function requires their optimal tuning.

### Adult-born immature GC biophysics prioritise energy economy along the Pareto front

The dentate gyrus is one of the few brain regions that undergoes adult neurogenesis (Altman and Das, 1965; Kuhn et al., 1996). Immature GCs play an important role in hippocampal memory formation (Clelland *et al*., 2009; Johnston et al., 2016), and contribute to pattern separation (Clelland *et al*., 2009; Sahay et al., 2011; Kropff et al., 2015; Tuncdemir et al., 2019; Yang *et al*., 2024).

To determine how these newly generated, immature GCs balance the computation-economy trade-off, we compared how well immature and mature GC models perform for the two objectives of energy economy and pattern separation in our population of GC models. Immature GCs are highly dynamic in their dendritic structure during maturation, but between 4 − 6 weeks after birth the total dendritic length stabilizes and the cells show increased synaptic plasticity (Ge *et al*., 2007; Kim et al., 2012). To specifically focus on biophysical synaptic and intrinsic properties, we modelled immature GCs at that stage of development, where they are morphologically similar to mature GCs but receive weaker and less specific synaptic inputs, and have increased intrinsic excitability due to reduced membrane conductances (for *Kir*2.1, *K*_*v*_2.1, *K*_*v*_4.2, *K*_*v*_7.2*/*3, *Ca*_*v*_1.3, *na*8*st, BK*, Mongiat *et al*., 2009; Beining et al., 2017b). To computationally represent distinct biophysics of mature and immature cells, we took a common set of GC morphologies (**Fig 5A**, Schmidt-Hieber *et al*., 2007) and adjusted their synaptic inputs and ionic conductances based on experimental knowledge, as we did previously (Beining *et al*., 2017b). Conspicuously, in the resulting function space, the mature GCs (*Blue shades*) displayed higher pattern separation while the immature GCs prioritised higher economy (*Red shades*, **Fig 5B**).

**Figure 5.**
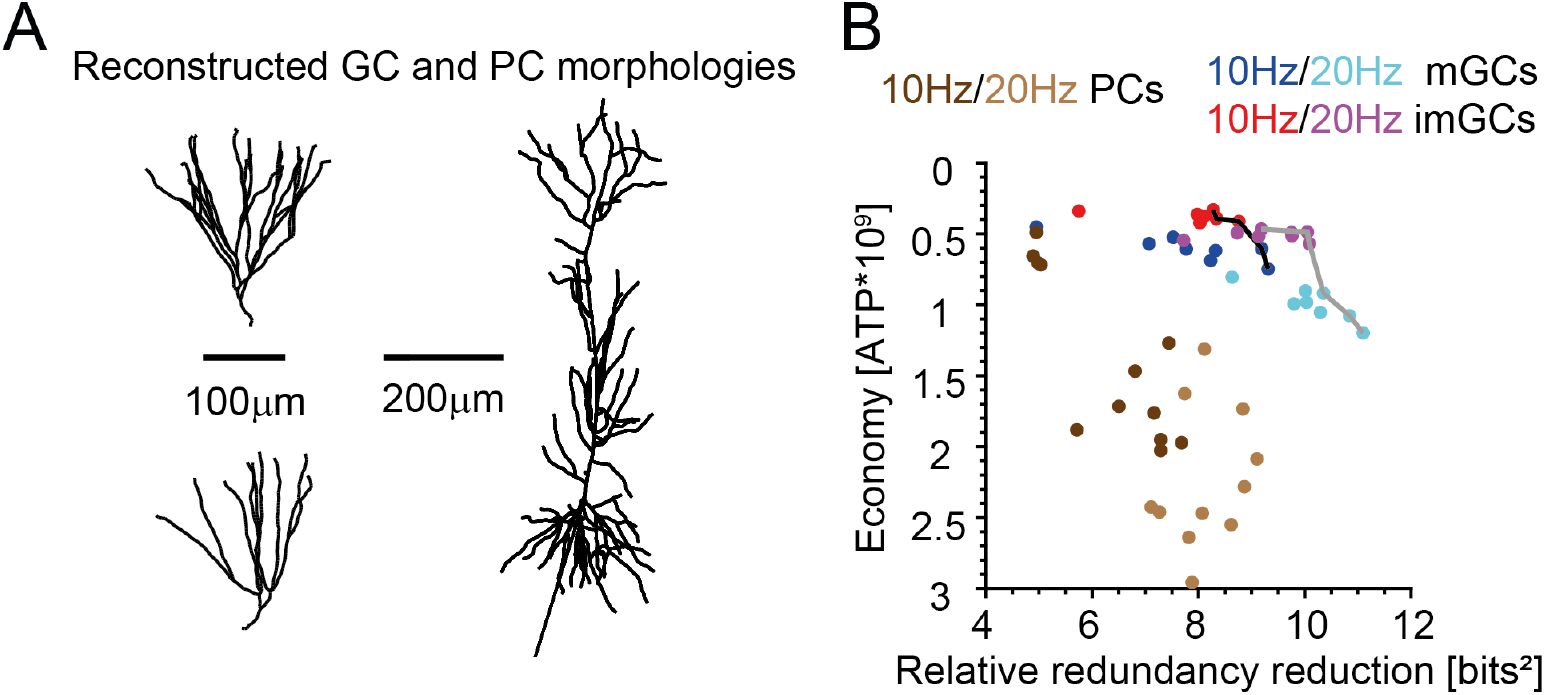
Mature and immature dentate GCs outperform CA1 pyramidal cells for both objectives with immature GCs prioritising metabolic efficiency over pattern separation. **A**: Example morphologies for deployed PCs and GCs. Diameter neglected for better visualisation. **B**: Economy vs. pattern separation shown for mature GCs (mGCs, *blue*), immature GCs (imGCs, *red*) and PCs (*brown*). The 20*Hz* condition is shown by the light colours. Pareto fronts are shown in black (10*Hz*) and grey (20*Hz*).

As a point of contrast, we evaluated the performance of standard hippocampal pyramidal cell (PC) models in the same function space. Hippocampal CA1 PCs are larger than GCs (**Fig 5A**) and are not typically associated with pattern separation (Jarsky *et al*., 2005; Hainmueller and Bartos, 2018). In agreement with this, their joint performance for both objectives was significantly worse than in both populations of GC models (**Fig 5B**, *brown*), confirming that CA1 PCs optimise different functional trade-offs than GCs.

What is the predominant mechanism behind the lower energy expenditure in immature GCs as compared with mature GCs both in our models and in real neurons? The weaker synaptic input is a major driver of this higher energy economy, as synaptic currents typically make up the majority of a neuron’s energetic demands (Attwell and Laughlin, 2001; Howarth et *al*., 2012). To maintain similar firing rates to the mature GCs despite the weaker synaptic drive, immature GCs compensate via reduced potassium conductance that increases their input resistance and intrinsic excitability. Interestingly, a similar mechanism of weakening synaptic inputs whilst increasing excitability to increase economy at the cost of computational performance was also recently seen in energy-restricted visual cortical neurons (Padamsey *et al*., 2022; Sommer et al., 2026).

To investigate this effect more thoroughly, we systematically altered the conductance of the *Kir*2.1 potassium channel, which is known to differ greatly between mature and immature GCs (in our model it was downregulated by 73%). Decreasing *Kir*2.1 channel density did not strongly impair the pattern separation performance of mature GCs (*blue*), but increasing it did (**Fig 6A**). In contrast, if the immature GCs (*red*) had the same channel density as the mature GCs, they would be severely impaired for pattern separation due to their reduced synaptic inputs (**Fig 6A**, Scaling of 1). The *Kir*2.1 channel therefore plays a major role in the pattern separation performance of the GCs. This is in line with previous experimental (Young *et al*., 2009; Pignatelli et al., 2019) and computational studies by us (Bird *et al*., 2024) and others (Yim *et al*., 2015). The developmental downregulation of the *Kir*2.1 channel prevents the severe disruption of pattern separation performance in the immature GC.

**Figure 6.**
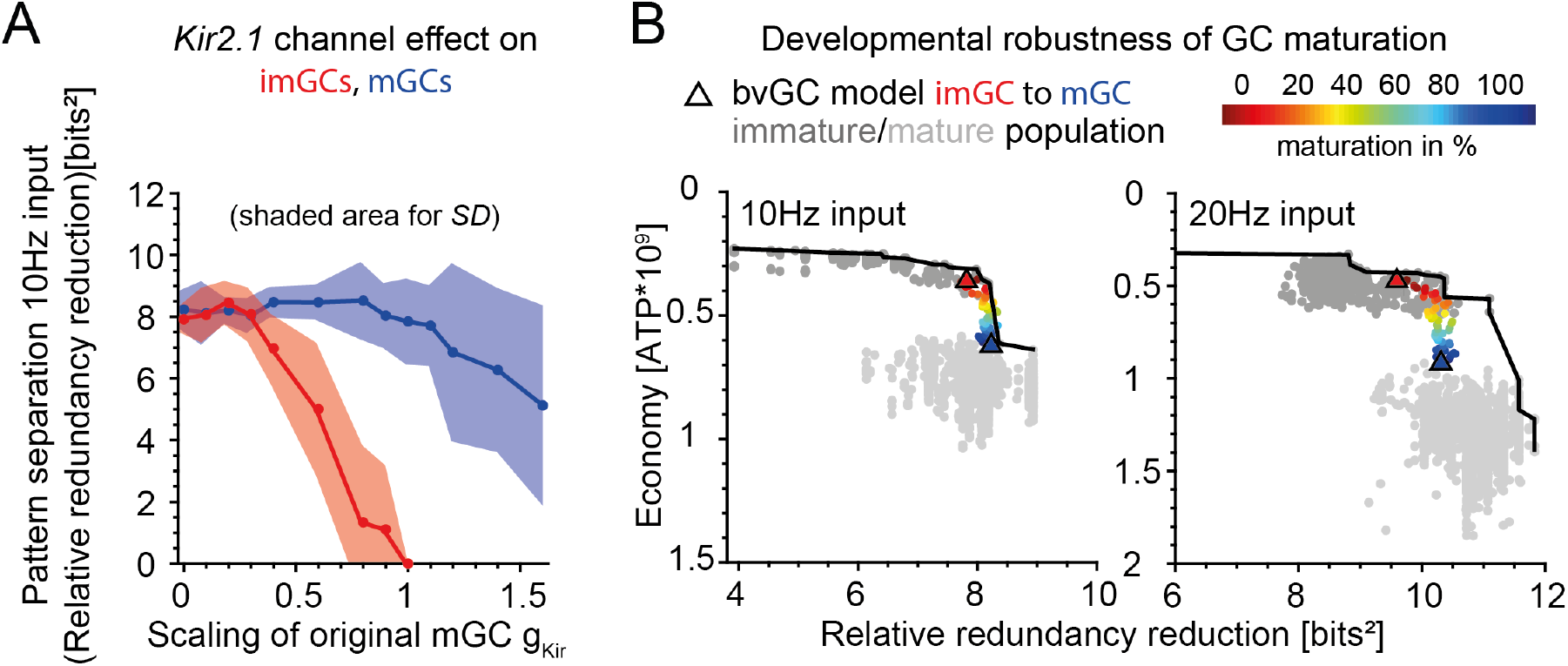
GCs move along the Pareto front during incremental in silico maturation. **A**: Influence of the *Kir*2.1 channel via conductance scaling on pattern separation performance for mature GCs (mGCs, *blue*) and immature GCs (imGCs, *red*); shaded area for standard deviation. **B**: Mean performance during GC maturation (*coloured dots/triangle*: bvGC model) compared to mature/immature population models (*light grey/ dark grey*). Respective Pareto lines are shown in black.

We then sought to explore the developmental trajectories of immature GCs in the two-objective performance space. By combining all known intrinsic (channel densities) and extrinsic (synaptic drive) changes of GCs during maturation (incremental steps of 2% from immature GC to mature GCs; 0% maturation *red triangle* and 100% maturation *blue triangle*), we investigated the robustness of their performance as they mature (**Fig 6B**, colour: mean performance of 8 cell morphologies from a data set of Schmidt-Hieber *et al*., 2007) with implemented biophysics from our bvGC model (Beining *et al*., 2017b). Remarkably, for the 10*Hz* input rate the developmental performance trajectory of the bvGC model follows the Pareto front (*Black line*) of optimal models from the immature (*dark grey*) to the mature (*light grey*) GC populations. This indicates that GCs remain Pareto optimal for pattern separation and energy consumption across development.

## Discussion

We have shown that dentate gyrus GCs are Pareto optimal for the performance of temporal pattern separation (Sahay *et al*., 2011; Schmidt et al., 2012; Guzman et al., 2021; Rolls, 2021; Yang *et al*., 2024) and the economy of energy consumption (Attwell and Laughlin, 2001; Quintela-López *et al*., 2022). This represents striking evidence that computational power and energetic constraints must be thought of together to truly understand a neuron’s biophysical principles (Remme *et al*., 2018; Fardet and Levina, 2020; Laughlin and Sejnowski, 2003). Throughout our investigation, Pareto optimality has been an invaluable guide, since improvement in computational performance typically comes with a trade-off in energetic cost. Both the bvGC model and the patch-clamped GC data had poorer pattern separation performance than some models in the degenerate population, but cells that exceeded experimental performance were notably more energetically costly. Other studies have observed a trade-off between energy consumption and the computational functions of transmitting visual information (Harris *et al*., 2015) or coincidence detection (Remme *et al*., 2018). It was not obvious that a similar trade-off should also exist for pattern separation, as sparse firing can theoretically be beneficial for both computation and its energy cost (Attwell and Laughlin, 2001; Cayco-Gajic and Silver, 2019; Bird *et al*., 2024). Nevertheless, we found that the two objectives are opposed for realistic cell models and inputs. Our simulations showed that the bvGC model, based on experimentally recorded ion channel densities throughout the dendrites (Beining *et al*., 2017b), consistently outperformed degenerate models fitted only to voltage traces, indicating that the biophysics of GCs are indeed highly specialised and support effective joint performance of these two functions.

The population of GC models, which was degenerate in terms of voltage response to somatic currents (Schneider *et al*., 2023), was surprisingly diverse in terms of both pattern separation and economy. When these functional tasks were incorporated into the assessment of the GCs, only a handful of models appeared viable. In general, it is not truly known what the total objective function of the GC is. GCs are involved in pattern separation (Yassa and Stark, 2011) and all cells must use their metabolic resources rationally (Alle *et al*., 2009; Sengupta *et al*., 2010; Schmidt-Hieber and Bischofberger, 2010; Yu and Yu, 2017). Our work strongly indicates that these two objectives are dominant in determining the observed GC biophysics. Other constraints or tasks could very well also be involved in the natural function of the GC, particularly beyond the single cell level. Our own results suggest that one such additional constraint is robustness to a range of input activity. Our relaxation of the voltage-degeneracy condition on the model population revealed that spiking models with the same larder of ion channels to choose from could functionally outperform real cells for specific inputs, but lost the ability of real cells to perform Pareto optimally across a range of realistic input patterns. One could also consider the function of pattern completion for mature GCs (Nakashiba *et al*., 2012b), other dendritic computations (London and Häusser, 2005; Larkum et al., 2007), or several plasticity phenomena (Morrison *et al*., 2008; Magee and Grienberger, 2020; Inglebert et *al*., 2020). Combining GC subpopulations (Yang *et al*., 2024; Mishra and Narayanan, 2019) at the network level might also reveal further constraints on their biophysics.

Degeneracy in general can be useful for the robustness of a system and its ability to attain and maintain stable performance (Whitacre and Bender, 2010). For example, failure in a particular ion channel might be compensated for by other degenerate channels (many-to-one mapping of channels to function, Rich *et al*., 2022; Schneider et al., 2023; Stö ber et al., 2023). Compensation for one task could at the same time perturb another, since the effect of an ion channel is usually not limited to one feature (see pleiotropy in Edelman and Gally, 2001; Yang and Prescott, 2023). Schneider et al. (2023) showed that degeneracy in spiking responses made it easier to identify valid neuron models and might contribute to the evolvability of neural systems (Whitacre and Bender, 2010). Adding functional constraints such as computational performance and energetic economy drastically reduced the degenerate parameter space of realistic models to those on the Pareto front. This highlights the degree of degeneracy that is necessary for neurons that are optimal for multiple realistic objectives (Stemmler and Koch, 1999; Jedlicka et al., 2022).

Although immature adult-born GCs contribute to pattern separation (Clelland *et al*., 2009; Sahay *et al*., 2011; Kropff et al., 2015; Tuncdemir et al., 2019; Yang et al., 2024), the computational division of labor and the interplay between immature and mature subpopulations is not fully understood (Kempermann, 2002; Jessberger and Kempermann, 2003; Lopez-Rojas and Kreutz, 2016; Johnston et al., 2016). We demonstrate that the specific biophysical properties of immature GCs maintain them on the Pareto front. However, although they remain Pareto optimal, they adopt a differently weighted trade-off, prioritizing lower energy consumption over higher pattern separation performance compared to mature GCs. While neurogenesis itself is an energy-intense process (Perera *et al*., 2011), the addition of a highly excitable and plastic cell population to the dentate gyrus network provides computational benefits, such as enhancing memory resolution (Aimone *et al*., 2011) or preventing catastrophic interference (Appleby and Wiskott, 2009). Because newborn immature GCs display increased responsiveness and plasticity (Schmidt-Hieber *et al*., 2004; Ge et al., 2007; Kim et al., 2012; Kropff et al., 2015; Kempermann, 2022) in combination with sparse connectivity (Lodge and Bischofberger, 2019), which benefits learning new information (Marín-Burgin and Schinder, 2012), their metabolic efficiency likely ensures that their activity and synaptic changes are energetically cheaper as they learn. The developmental trajectories we have simulated using stepwise biophysical maturation informed by experimental data (Mongiat *et al*., 2009; Lodge and Bischofberger, 2019) indicate that maturing GCs move along the Pareto front, continuously adjusting the balance between economy and function while remaining optimal for these two objectives at all stages.

## Materials and Methods

### Software

Morphological analysis was performed in Matlab (Mathworks Inc, version 2018a and 2023b) using the software package of the `TREES toolbox` (Cuntz *et al*., 2011; www.treestoolbox.org). All passive and active compartmental model simulations were run in NEURON (Hines and Carnevale, 2004) via a newly developed software to control NEURON with Matlab and the TREES toolbox, the TREES-to-NEURON (`T2N`) interface (Beining *et al*., 2017b). The pattern separation measures were calculated with the pattern separation toolbox (Bird *et al*., 2024; www.github.com/synapsesanddendrites/Pattern_separation_toolbox).

### Morphological and compartmental models

The morphologies used in this research were reconstructed from 2 − 4 months old *C*57*BL/*6 mice (*n* = 8, Schmidt-Hieber *et al*., 2007). The compartmental model by Beining et al. (2017b) was implemented for the mature GCs. It consists of 16 voltage-gated ion channels including region- and voltage-dependent Na^+^, K^+^ and Ca^2+^ channels as well as calcium-dependent K^+^ channels (SK, BK); these channels typically have different densities in different dendritic layers.

Further, a realistic axon (350*µm*) was attached to the GC morphology and the eight-state sodium channel *na*8*st* within was adapted to correspond to energy data according to Alle *et al*. (2009), see **Figure 2B**. The parameters *ah*_0_ and *bh*_0_ were quadrupled to speed up inactivation. The performance was similar to the condition without axon (compare **Figure 3B** and **Figure S2**).

In order to simulate immature GCs, the same reconstructed morphologies as for the mature GCs were applied but with decreased synaptic density (30%) and (partially) blocked channels (*Kir*2.1, *K*_*v*_2.1, *K*_*v*_4.2, *K*_*v*_7.2*/*3, *Ca*_*v*_1.3, *na*8*st, BK*) according to literature (Mongiat *et al*., 2009; Beining *et al*., 2017b).

In comparison, a data set that included 12 mouse hippocampal CA1 PC morphologies from 10 − 14 months old WT mice was employed (Šišková *et al*., 2014). For the topological characteristics of the PC morphologies see previous work by Mittag et al. (2023). An artificial cosine shaped soma (surface area of 560*µm*^2^) was added to the dendrites as none of the reconstructions included an original soma (Benavides-Piccione *et al*., 2020). An artificial axon was created as well. It consisted of cylindrical compartments totalling a length of 630*µm* with an average diameter of 0.5*µm*. A biophysically realistic model based on the model developed by Poirazi et al. (2003b,a) was implemented for the PCs. The model consisted of active and passive membrane mechanisms including 16 different ion channels that were mostly heterogeneously distributed.

### Population models

The original GC model was used as the basis for a population of models to study the impact of ion channel parameters on GC function (Schneider *et al*., 2023). In total, 20, 000 GC models were generated by variation of 27 parameters, especially the channel conductances, in a range of [0, 2] times the original value. Approximately 7% of the model population were considered “good” regarding a *Fitness* function (*F <* 2) that calculated the closeness of the somatic electrophysiological behaviour to experiments.

### Excitatory synapses and their activation

In the dentate GC model by Beining *et al*. (2017b) excitatory, glutamate Alpha-amino-3-hydroxy-5-Methyl-4-isoxazole-Propionic Acid (AMPA) background synapses were implemented along the dendrites. The synapses were realised as time-dependent conductances following a dual-exponential with rise time constant of *τ*_1_ = 0.2*ms* and decay time constant of *τ*_2_ = 2.5*ms* (*`Exp2Syn`*NEURON object). The reversal potential was 0*mV* . N-Methyl-DAspartate (NMDA) synapses were included with a voltage-dependence that is non-linear. They consisted of a magnesium block which could be released with strong enough depolarisation. Compared to AMPA synapses the NMDA synapses had slower exponentials with rise time constant of *τ*_1_ = 0.33*ms* and decay time constant of *τ*_2_ = 50*ms* with the same reversal potential of 0*mV* (*`Exp2nmda2`*NEURON object based on Krueppel *et al*., 2011). The NMDA synapses matched the densities of the AMPA synapses and were excited by the same presynaptic Poisson input. Realistic, inhomogeneous synaptic distributions for the densities of the AMPA and NMDA synapses were implemented (Schmidt-Hieber *et al*., 2007).

**Table 1.** Dentate Gyrus GC synapse densities.

| Dendritic region | GCL | IML | MML | OML |
| --- | --- | --- | --- | --- |
| AMPA, NMDA synapse/$\mu m$ | 0.05 | 0.1 | 0.2 | 0.05 |

Activation of the background synapses was implemented via a *`VecStim`*point processes with Poisson spike trains with an average spiking frequency that ranged from 1*Hz* to 2*Hz* (T2N function *`t2n poissonSpikeGen`*).

Perforant pathway input (from EC) to the GCs was implemented as the main source of patterns to separate. In order to create an ensemble of spike trains with specific train number, Poisson frequency, phase and correlation the `pattern separation toolbox` was used (Bird *et al*., 2024). This was necessary to compare input (6 random key input synapses in the outer molecular layer) and output (evaluated at the AIS) spike trains for the pattern separation measure as described below. The detailed activation frequencies of the experiments are mentioned in the corresponding section.

### Simulation setup

Known input frequencies of GCs in DG projecting from the EC were used (the perforant pathway with *theta* and *beta* signals between 5*Hz* and 20*Hz*, Quintanilla *et al*., 2022). In detail, three different frequencies with three different spike train patterns respectively were successively fed into six key input synapses of the GC outer molecular layer: phase-locked spike ensemble with three component trains, a spiking rate of 10*Hz*, 15*Hz* and 20*Hz*, a correlation strength of 0.7 and a phase rate of 6*Hz*. By comparing input and output spike patterns via the `pattern separation toolbox` by Bird et al. (2024) the relative redundancy reduction and sparsity weighted MI were calculated as in **Equations 15** and **17** to get the pattern separation measure.

During the simulation time of 2*s* the ATP consumption (energy used for the extrusion of sodium and calcium as in **Equation 11**) with contributions from the channels and synapses (**Equation 7**) was recorded. For all valid GC models (Schneider *et al*., 2023) the economy of energy and the pattern separation performance were calculated and compared. A model was decided to be on the Pareto front if it could not be outperformed by any other model, in both economy and pattern separation, at the same time.

### ATP consumption

In order to calculate the energy of the cell activity an ion counting system was used (Attwell and Laughlin, 2001). The Na^+^ ions needed for the postsynaptic currents and APs are the main contributors to the energy budget (Attwell and Laughlin, 2001). For example, after a concentration change due to the flux of Na^+^ and K^+^ for an AP generation, the ion concentrations have to be restored again by ion pumps. In general, the Na^+^*/*K^+^ pump extrudes three Na^+^ and imports two K^+^ ions per one adenosine triphosphate (*ATP* ) molecule (Post and Jolly, 1957). Therefore, for the calculation of the energy the *ATP* consumption, we used with the following relations, similar to Remme et al. (2018):

- Na^+^*/*K^+^ pump with 1*ATP* consumption ≡ 3 Na^+^ ions extrusion.
- Na^+^*/*Ca^2+^ exchange with 1 Ca^2+^ ion ≡ 3 Na^+^ ions.
- Charge *Q* of 1*C* of Na^+^ with 6.24 *·* 10^18^ particles.
- Charge *Q* of 2*C* of Ca^2+^ with 6.24 *·* 10^18^ particles.

All currents of the *I*_*Ca*_ channels were read out directly with the code. The sodium current however consisted of several contributions:

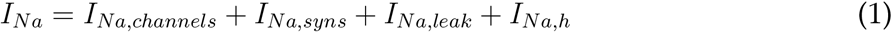

For the first term, the sodium channels were read out with the code. The other three terms were determined by the following calculations, e.g. synaptic transmission:

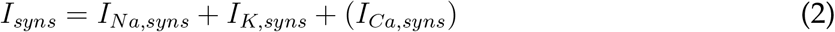

The synaptic transmission included cations only. In AMPA synapses the Ca^2+^ ions were neglected (see Harris *et al*., 2015).

Similarly, the Leak current consisted of several contributions:

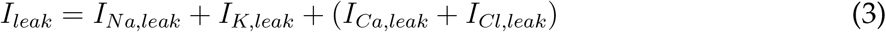

External Ca^2+^ was neglected as it only indirectly influences the leak sodium channel (see Lu *et al*., 2010). Chloride was neglected as well since its influence occurs via GABA synapses which were not included in the set up. With Ohms law, the following expression was derived:

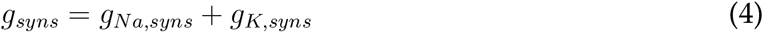

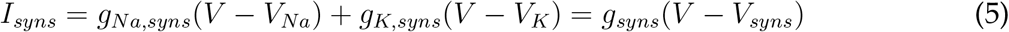

With the equilibrium potential for the synapses at *V*_*syns*_ = 0. Setting *V* = 0:

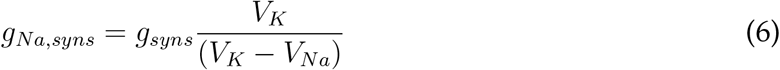

It follows for *I*_*Na,syns*_:

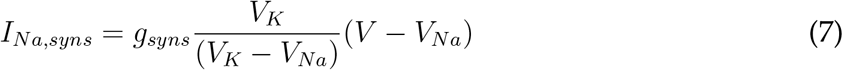

With *g*_*syns*_ = *g*_*AMPA*_.

Under the assumption of minimal Ca^2+^ entry through NMDA, the same expression for *I*_*Na,NMDA*_ with *g*_*syns*_ = *g*_*NMDA*_ follows.

The same derivation as for *I*_*Na,syns*_ was done for the sodium contributions of *I*_*Na,leak*_ and *I*_*Na,h*_ (leak and HCN channel):

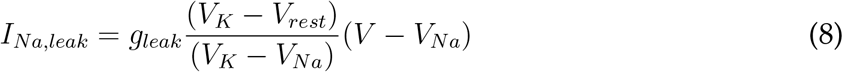

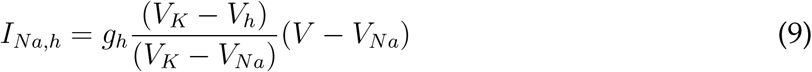

With the equilibrium potentials at *V*_*Na*_ = 87.76*mV, V*_*K*_ = −93*mV* and *V*_*h*_ = −41.9*mV* respectively.

With the total sodium current *I*_*Na*_ and calcium current *I*_*Ca*_ per time step, consequently, the time dependent integral over the whole stimulation period *T* was computed to get the charge:

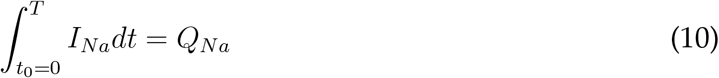

The same goes for *Q*_*Ca*_.

With the relations above, total *ATP* was calculated as *ATP* = *ATP*_*Na*_ + *ATP*_*Ca*_. One charge of sodium relates to 1*/*3 *·* 6.24 *·* 10^18^ *ATP* molecules and two charge of calcium relate to 6.24 *·* 10^18^ *ATP* molecules, yielding:

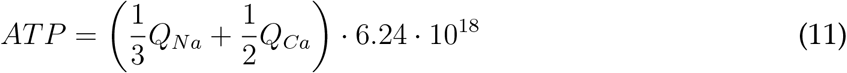

### Pattern separation and mutual information

According to Shannon (1948), the expected information *I*(*X*) in *bits* of a random variable *X* can also be called uncertainty or entropy *H*(*X*). It is dependent on the probability *p* (between 0 and 1) of its potential values and defined in the following:

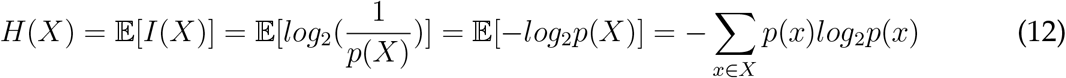

Furthermore, the MI of *I*(*X, Y* ) between variables *X* and *Y* measures the decrease of uncertainty over *Y* by knowing about *X* or vice versa (Shannon, 1948):

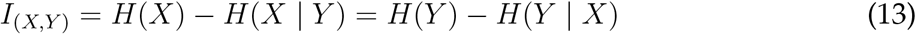

With *H*(*X* | *Y* ) being the conditional entropy and with Bayes’ rule this yields:

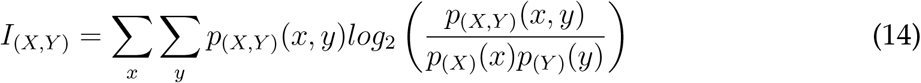

Here, *p*(*X, Y* ), *p*(*X*) and *p*(*Y* ) are the joint and marginal distributions respectively. For independent *X* and *Y, I*_(*X,Y* )_ equals 0. This can be applied to neural spike trains with input and output patterns *X* and *Y* . For a combined measure of pattern separation and information theory (Bird *et al*., 2024) this MI was weighted with the number of spikes *m*_*x*_ and *n*_*y*_ of *X* and *Y* to give:

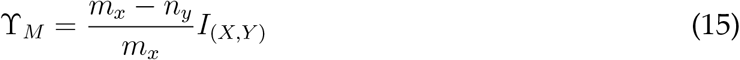

Next, the redundancy *R*_*X*_ within input spike trains *X* was considered as the minimum MI between any part or single train *x*_*i*_ and the rest of the ensemble *X\x*_*i*_ (Williams and Beer, 2010):

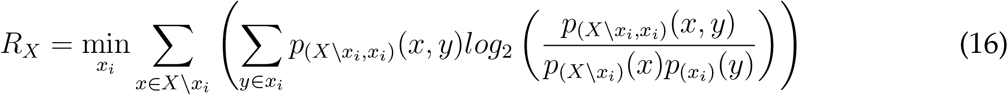

Similarly, *R*_*Y*_ was calculated for the output spike trains *Y* . Relative redundancy reduction as a new measure for pattern separation then compared both redundancies and weighed them with the MI of *I*(*X, Y* ):

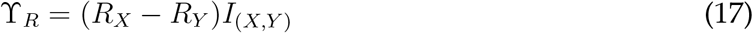

## Acknowledgments

We acknowledge funding through DFG 467764793, JE 528/10-1. We would like to thank Marcel Beining for helpful discussions on this work.

## Supplement

**Figure S1.**
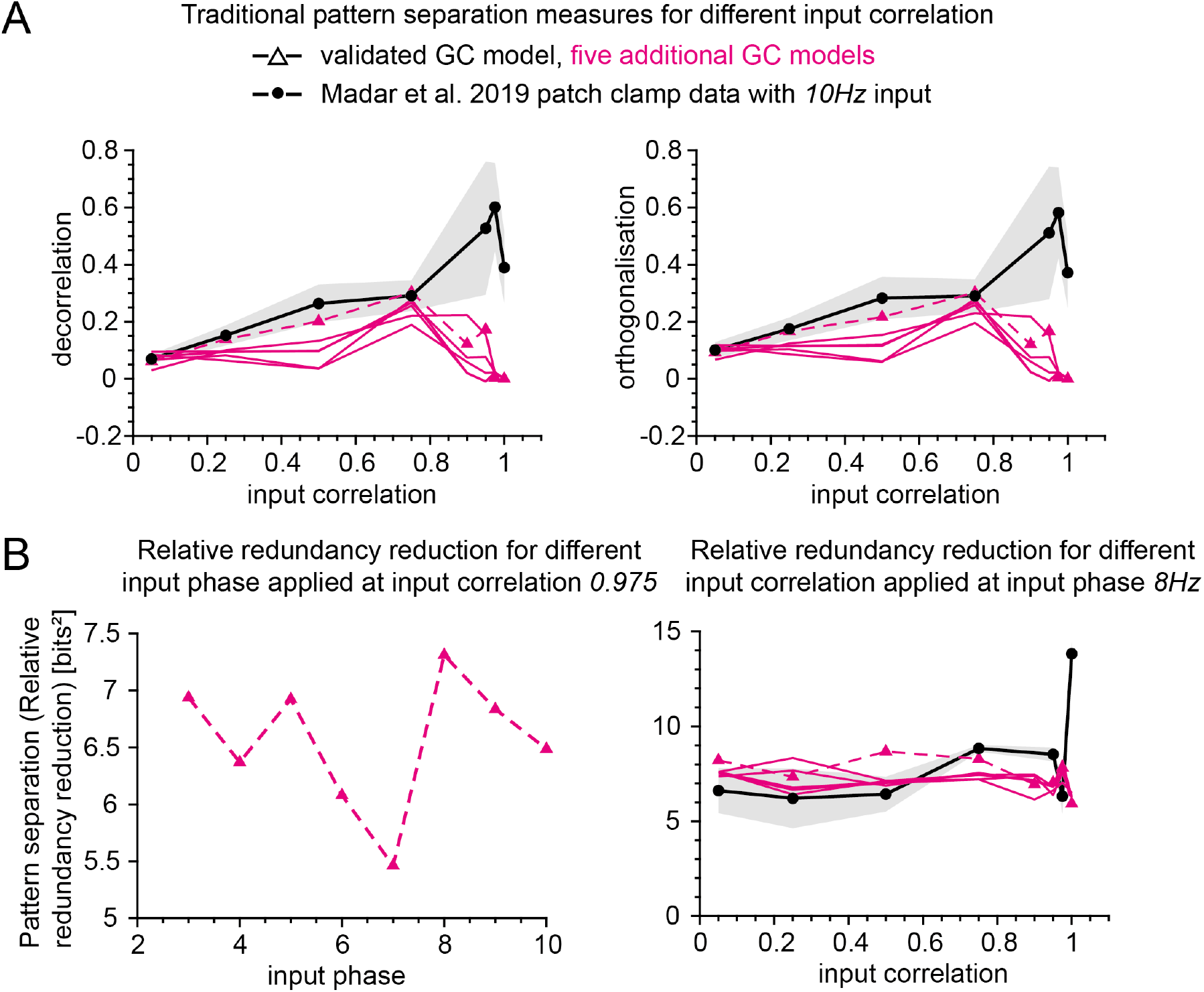
Additional pattern separation measures compared to Madar *et al*. (2019) **A**: Traditional pattern separation measures (orthogonalisation and decorrelation) of Madar et al. (2019) and simulated GCs diverge. **B**: Effect of input phase on relative redundancy reduction (left: for various input phase and constant correlation 0.975; right: for fixed input phase of 8Hz and various correlation).

**Table S1.** Sodium per AP in various nerve cells: mossy fiber bouton MFB as in Alle et al. (2009); models of squid giant axon SA, crab motor neuron axon CA, mouse fast-spiking neuron MFS, honeybee Kenyon cell BK, rat hippocampal interneuron RHI, rat granule cell RGC, mouse thalamo-cortical relay neuron MTCR as in Sengupta et al. (2010); and data from CA1 pyramidal cell PC, cortical interneuron CIN, purkinje cell PuC and cortical pyramidal cell CPC from Carter and Bean (2009). Sodium overlap/non-overlap means that the sodium is/is not counterbalanced by simultaneous flowing of potassium. All currents are in [*nC/cm*^2^].

| Nerve Cell | Sodium total | Capacitive minimum | Sodium non-overlap | Sodium overlap | Non-overlap /Total [%] | Capacitive /Total [%] |
| --- | --- | --- | --- | --- | --- | --- |
| MFB | 153 | 121 | 122.4 | 30.6 | 80 | 79 |
| SA | 1098 | 98 | 64 | 1034 | 5.8 | 8.9 |
| CA | 364 | 106 | 100 | 264 | 27.47 | 29.12 |
| MFS | 315 | 129 | 84 | 231 | 26.7 | 41 |
| BK | 186 | 61 | 57 | 129 | 30.6 | 32.8 |
| RHI | 76 | 56 | 40 | 36 | 52.6 | 73.7 |
| RGC | 72 | 69 | 63 | 9 | 87.5 | 95.8 |
| MTCR | 65 | 65 | 50 | 15 | 76.9 | 100 |
| CA1 PC | 122.55 | 75.65 | 75.65 | 46.9 | 61.73 | 61.73 |
| CIN | 131.24 | 66.28 | 66.28 | 64.96 | 50.50 | 50.50 |
| PuC | 148.61 | 74.31 | 74.31 | 74.3 | 50 | 50 |
| CPC | 79.13 | 63.81 | 63.81 | 15.32 | 80.64 | 80.64 |

**Figure S2.**
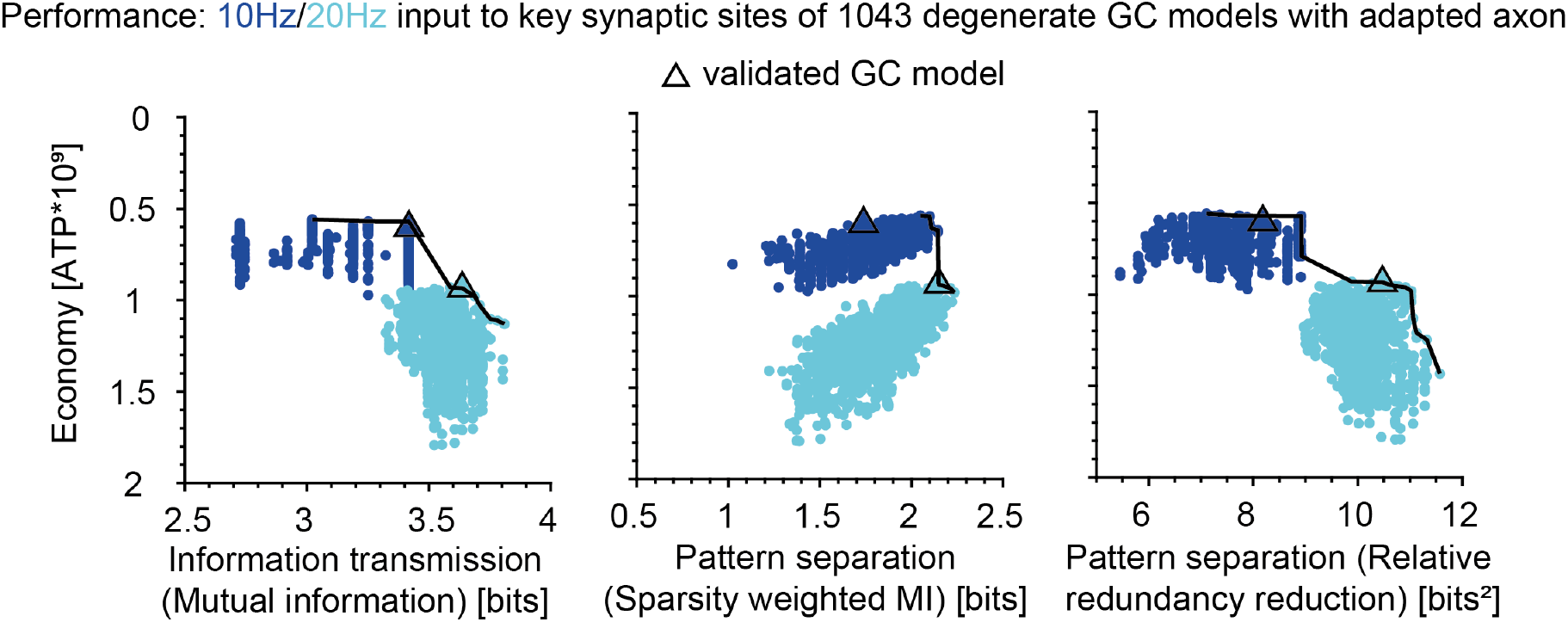
Economy and pattern separation in GC model population with adapted axon.

## References

Aimone JB, Deng W, Gage FH (2011) Resolving new memories: a critical look at the dentate gyrus, adult neurogenesis, and pattern separation. Neuron 70:589–596.

Albus J (1971) A theory of cerebellar function. Math. Biosci. 10:25–61.

Alle H, Roth A, Geiger JR (2009) Energy-efficient action potentials in hippocampal mossy fibers. Science 325:1405–1408.

Alon U (2006) An introduction to systems biology: design principles of biological circuits Chapman and Hall/CRC.

Altman J, Das GD (1965) Autoradiographic and histological evidence of postnatal hippocampal neurogenesis in rats. J Comp Neurol 124:319–335.

Appleby PA, Wiskott L (2009) Additive neurogenesis as a strategy for avoiding interference in a sparsely-coding dentate gyrus. Network: Computation in Neural Systems 20:137–161.

Attwell D, Laughlin SB (2001) An energy budget for signaling in the grey matter of the brain. Journal of Cerebral Blood Flow & Metabolism 21:1133–1145.

Bakker A, Kirwan CB, Miller M, Stark CE (2008) Pattern separation in the human hippocampal ca3 and dentate gyrus. science 319:1640–1642.

Bast A, Fruengel R, de Kock CP, Oberlaender M (2023) Interpretable modelling of input-output computations in cortex. bioRxiv pp. 2023–08.

Beining M, Jungenitz T, Radic T, Deller T, Cuntz H, Jedlicka P, Schwarzacher SW (2017a) Adult-born dentate granule cells show a critical period of dendritic reorganization and are distinct from developmentally born cells. Brain Structure & Function 222:1427–1446.

Beining M, Mongiat LA, Schwarzacher SW, Cuntz H, Jedlicka P (2017b) T2N as a new tool for robust electrophysiological modeling demonstrated for mature and adult-born dentate granule cells. Elife 6:e26517.

Benavides-Piccione R, Regalado-Reyes M, Fernaud-Espinosa I, Kastanauskaite A, Tapia-González S, LeónEspinosa G, Rojo C, Insausti R, Segev I, Defelipe J (2020) Differential structure of hippocampal CA1 pyramidal neurons in the human and mouse. Cerebral Cortex 30:730–752.

Bird AD, Cuntz H, Jedlicka P (2024) Robust and consistent measures of pattern separation based on information theory and demonstrated in the dentate gyrus. PLoS Comput Biol 20:e1010706.

Carter BC, Bean BP (2009) Sodium entry during action potentials of mammalian neurons: incomplete inactivation and reduced metabolic efficiency in fast-spiking neurons. Neuron 64:898–909.

Cayco-Gajic N, Silver R (2019) Re-evaluating circuit mechanisms underlying pattern separation. Neuron 101:584–602.

Chavlis S, Poirazi P (2017) Pattern separation in the hippocampus through the eyes of computational modeling. Synapse 71:e21972.

Clelland C, Choi M, Romberg C, Clemenson Jr G, Fragniere A, Tyers P, Jessberger S, Saksida L, Barker R, Gage F et al. (2009) A functional role for adult hippocampal neurogenesis in spatial pattern separation. Science 325:210–213.

Crotty P, Levy WB (2007) Effects of na+ channel inactivation kinetics on metabolic energy costs of action potentials. Neurocomputing 70:1652–1656.

Cuntz H, Forstner F, Borst A, Häusser M (2011) The TREES Toolbox Probing the basis of axonal and dendritic branching. Neuroinformatics 9:91–96.

Danielson NB, Kaifosh P, Zaremba JD, Lovett-Barron M, Tsai J, Denny CA, Balough EM, Goldberg AR, Drew LJ, Hen R, Losonczy A, Kheirbek MA (2016) Distinct contribution of adult-born hippocampal granule cells to context encoding. Neuron 90:101–112.

Edelman GM, Gally JA (2001) Degeneracy and complexity in biological systems. Proceedings of the National Academy of Sciences of the United States of America 98:13763–13768.

Fardet T, Levina A (2020) Simple models including energy and spike constraints reproduce complex activity patterns and metabolic disruptions. PLoS Computational Biology 16:e1008503.

Finnegan R, Becker S (2015) Neurogenesis paradoxically decreases both pattern separation and memory interference. Frontiers in Systems Neuroscience 9.

Ge S, Yang Ch, Hsu Ks, Ming Gl, Song H (2007) A critical period for enhanced synaptic plasticity in newly generated neurons of the adult brain. Neuron 54:559–566.

Groisman AI, Yang SM, Schinder AF (2020) Differential coupling of adult-born granule cells to parvalbumin and somatostatin interneurons. Cell Reports 30:202–214.e4.

Guzman SJ,Schlögl A, Espinoza C, Zhang X, Suter BA, Jonas P (2021) How connectivity rules and synaptic properties shape the efficacy of pattern separation in the entorhinal cortex–dentate gyrus–ca3 network. Nature Computational Science 1:830–842.

Hainmueller T, Bartos M (2018) Parallel emergence of stable and dynamic memory engrams in the hippocampus. Nature 558:292–296.

Harris JJ, Jolivet R, Engl E, Attwell D (2015) Energy-efficient information transfer by visual pathway synapses. Current Biology 25:3151–3160.

Hart Y, Sheftel H, Hausser J, Szekely P, Ben-Moshe NB, Korem Y, Tendler A, Mayo AE, Alon U (2015) Inferring biological tasks using pareto analysis of high-dimensional data. Nature methods 12:233–235.

Hines ML, Carnevale NT (2004) Discrete event simulation in the NEURON environment. Neurocomputing 58-60:1117–1122.

Hodgkin AL, Huxley AF, Katz B (1952) Measurement of current-voltage relations in the membrane of the giant axon of loligo. The Journal of physiology 116:424.

Howarth C, Gleeson P, Attwell D (2012) Updated energy budgets for neural computation in the neocortex and cerebellum. Journal of Cerebral Blood Flow & Metabolism 32:1222–1232.

Inglebert Y, Aljadeff J, Brunel N, Debanne D (2020) Synaptic plasticity rules with physiological calcium levels. Proceedings of the National Academy of Sciences 117:33639–33648.

Jarsky T, Roxin A, Kath WL, Spruston N (2005) Conditional dendritic spike propagation following distal synaptic activation of hippocampal CA1 pyramidal neurons. Nature Neuroscience 8:1667–1676.

Jedlicka P, Bird AD, Cuntz H (2022) Pareto optimality, economy–effectiveness trade-offs and ion channel degeneracy: improving population modelling for single neurons. Open Biology 12:220073.

Jessberger S, Kempermann G (2003) Adult-born hippocampal neurons mature into activity-dependent responsiveness. European Journal of Neuroscience 18:2707–2712.

Johnston ST, Shtrahman M, Parylak S, Gonçalves JT, Gage FH (2016) Paradox of pattern separation and adult neurogenesis: A dual role for new neurons balancing memory resolution and robustness. Neurobiology of learning and memory 129:60–68.

Kempermann G (2002) Why new neurons? possible functions for adult hippocampal neurogenesis. Journal of neuroscience 22:635–638.

Kempermann G (2022) What is adult hippocampal neurogenesis good for? Frontiers in Neuroscience 16:852680.

Kim WR, Christian K, Ming GL, Song H (2012) Time-dependent involvement of adult-born dentate granule cells in behavior. Behavioural brain research 227:470–479.

Kropff E, Yang SM, Schinder AF (2015) Dynamic role of adult-born dentate granule cells in memory processing. Current opinion in neurobiology 35:21–26.

Krueppel R, Remy S, Beck H (2011) Dendritic integration in hippocampal dentate granule cells. Neuron 71:512–528.

Kuhn H, Dickinson-Anson H, Gage F (1996) Neurogenesis in the dentate gyrus of the adult rat: Age-related decrease of neuronal progenitor proliferation. J Neurosci 16:2027–2033.

Larkum ME, Waters J, Sakmann B, Helmchen F (2007) Dendritic spikes in apical dendrites of neocortical layer 2/3 pyramidal neurons. Journal of Neuroscience 27:8999–9008.

Laughlin SB, Sejnowski TJ (2003) Communication in neuronal networks. Science 301:1870–1874.

Leutgeb JK, Leutgeb S, Moser MB, Moser EI (2007) Pattern separation in the dentate gyrus and ca3 of the hippocampus. science 315:961–966.

Lodge M, Bischofberger J (2019) Synaptic properties of newly generated granule cells support sparse coding in the adult hippocampus. Behavioural Brain Research 372:112036.

Löffler H, Gupta DS (2022) A Model of Pattern Separation by Single Neurons. Frontiers in Computational Neuroscience 16:858353.

London M, Häusser M (2005) Dendritic computation. Annu. Rev. Neurosci. 28:503–532.

Lopez-Rojas J, Kreutz MR (2016) Mature granule cells of the dentate gyrus—passive bystanders or principal performers in hippocampal function? Neuroscience & Biobehavioral Reviews 64:167–174.

Lu B, Zhang Q, Wang H, Wang Y, Nakayama M, Ren D (2010) Extracellular calcium controls background current and neuronal excitability via an unc79-unc80-nalcn cation channel complex. Neuron 68:488–499.

Madar AD, Ewell LA, Jones MV (2019) Pattern separation of spiketrains in hippocampal neurons. Scientific Reports 9:5282.

Magee JC, Grienberger C (2020) Synaptic plasticity forms and functions. Annual review of neuroscience 43:95–117.

Marr D (1971) Simple memory: A theory for archicortex. Phys. Trans. R. Soc. 262:23–81.

Marr D (1969) A theory of cerebellar cortex. Journal of Physiology pp. 437–470.

Marín-Burgin A, Schinder AF (2012) Requirement of adult-born neurons for hippocampus-dependent learning. Behavioural Brain Research 227:391–399.

Mishra P, Narayanan R (2019) Disparate forms of heterogeneities and interactions among them drive channel decorrelation in the dentate gyrus: Degeneracy and dominance. Hippocampus 29:378–403.

Mittag M, Mediavilla L, Remy S, Cuntz H, Jedlicka P (2023) Modelling the contributions to hyperexcitability in a mouse model of Alzheimer’s disease. The Journal of Physiology pp. 1–35.

Mongiat LA, Espósito MS, Lombardi G, Schinder AF (2009) Reliable activation of immature neurons in the adult hippocampus. PloS one 4:e5320.

Mongiat LA, Schinder AF (2011) Adult neurogenesis and the plasticity of the dentate gyrus network. European Journal of Neuroscience 33:1055–1061.

Morrison A, Diesmann M, Gerstner W (2008) Phenomenological models of synaptic plasticity based on spike timing. Biological cybernetics 98:459–478.

Nakashiba T, Cushman JD, Pelkey KA, Renaudineau S, Buhl DL, McHugh TJ, Barrera VR, Chittajallu R, Iwamoto KS, McBain CJ, Fanselow MS, Tonegawa S (2012a) Young dentate granule cells mediate pattern separation whereas old granule cells contribute to pattern completion. Cell 149:188–201.

Nakashiba T, Cushman JD, Pelkey KA, Renaudineau S, Buhl DL, McHugh TJ, Barrera VR, Chittajallu R, Iwamoto KS, McBain CJ et al. (2012b) Young dentate granule cells mediate pattern separation, whereas old granule cells facilitate pattern completion. Cell 149:188–201.

Neunuebel JP, Knierim JJ (2014) Ca3 retrieves coherent representations from degraded input: direct evidence for ca3 pattern completion and dentate gyrus pattern separation. Neuron 81:416–427.

Niven JE (2016) Neuronal energy consumption: biophysics, efficiency and evolution. Current opinion in neurobiology 41:129–135.

Padamsey Z, Katsanevaki D, Dupuy N, Rochefort NL (2022) Neocortex saves energy by reducing coding precision during food scarcity. Neuron 110:280–296.

Pallasdies F, Norton P, Schleimer JH, Schreiber S (2021) Neural optimization: Understanding trade-offs with pareto theory. Current Opinion in Neurobiology 71:84–91.

Perera TD, Lu D, Thirumangalakudi L, Smith EL, Yaretskiy A, Rosenblum LA, Kral JG, Coplan JD et al. (2011) Correlations between hippocampal neurogenesis and metabolic indices in adult nonhuman primates. Neural plasticity 2011.

Pignatelli M, Ryan TJ, Roy DS, Lovett C, Smith LM, Muralidhar S, Tonegawa S (2019) Engram cell excitability state determines the efficacy of memory retrieval. Neuron 101:274–284.

Poirazi P, Brannon T, Mel BW (2003a) Arithmetic of subthreshold synaptic summation in a model CA1 pyramidal cell. Neuron 37:977–987.

Poirazi P, Brannon T, Mel BW (2003b) Pyramidal neuron as two-layer neural network. Neuron 37:989–999.

Post RL, Jolly PC (1957) The linkage of sodium, potassium, and ammonium active transport across the human erythrocyte membrane. Biochimica et biophysica acta 25:118–128.

Quintanilla J, Jia Y, Lauterborn JC, Pruess BS, Le AA, Cox CD, Gall CM, Lynch G, Gunn BG (2022) Novel types of frequency filtering in the lateral perforant path projections to dentate gyrus. The Journal of Physiology 600:3865–3896.

Quintela-López T, Shiina H, Attwell D (2022) Neuronal energy use and brain evolution. Current Biology 32:R650–R655.

Remme MW, Rinzel J, Schreiber S (2018) Function and energy consumption constrain neuronal biophysics in a canonical computation: Coincidence detection. PLoS computational biology 14:e1006612.

Restivo L, Niibori Y, Mercaldo V, Josselyn SA, Frankland PW (2015) Development of Adult-Generated Cell Connectivity with Excitatory and Inhibitory Cell Populations in the Hippocampus. Journal of Neuroscience 35:10600–10612.

Rich S, Chameh HM, Lefebvre J, Valiante TA (2022) Loss of neuronal heterogeneity in epileptogenic human tissue impairs network resilience to sudden changes in synchrony. Cell Reports 39:110863.

Rolls ET (2021) On pattern separation in the primate, including human, hippocampus. Trends in Cognitive Sciences 25:920–922.

Sahay A, Scobie KN, Hill AS, O’Carroll CM, Kheirbek MA, Burghardt NS, Fenton AA, Dranovsky A, Hen R (2011) Increasing adult hippocampal neurogenesis is sufficient to improve pattern separation. Nature 472:466–470.

Sahay A, Wilson D, Hen R (2011) Pattern separation: A common function for new neurons in hippocampus and olfactory bulb. Neuron 70.

Schmidt B, Marrone DF, Markus EJ (2012) Disambiguating the similar: the dentate gyrus and pattern separation. Behavioural brain research 226:56–65.

Schmidt-Hieber C, Bischofberger J (2010) Fast sodium channel gating supports localized and efficient axonal action potential initiation. Journal of Neuroscience 30:10233–10242.

Schmidt-Hieber C, Jonas P, Bischofberger J (2004) Enhanced synaptic plasticity in newly generated granule cells of the adult hippocampus. Nature 429:184–187.

Schmidt-Hieber C, Jonas P, Bischofberger J (2007) Subthreshold dendritic signal processing and coincidence detection in dentate gyrus granule cells. Journal of Neuroscience 27:8430–8441.

Schneider M, Bird AD, Gidon A, Triesch J, Jedlicka P, Cuntz H (2023) Biological complexity facilitates tuning of the neuronal parameter space. PLOS Computational Biology 19:e1011212.

Sengupta B, Stemmler M, Laughlin SB, Niven JE (2010) Action potential energy efficiency varies among neuron types in vertebrates and invertebrates. PLoS computational biology 6:e1000840.

Shannon CE (1948) A mathematical theory of communication. The Bell system technical journal 27:379–423.

Sheftel H, Shoval O, Mayo A, Alon U (2013) The geometry of the p areto front in biological phenotype space. Ecology and evolution 3:1471–1483.

Šišková Z, Justus D, Kaneko H, Friedrichs D, Henneberg N, Beutel T, Pitsch J, Schoch S, Becker A, von der Kammer H, Remy S (2014) Dendritic structural degeneration is functionally linked to cellular hyperexcitability in a mouse model of Alzheimer’s disease. Neuron 84:1023–1033.

Sommer P, Zeldenrust F, Jedlicka P, Bird AD, Triesch J (2026) Neuronal degeneracy reflects context-specific trade-offs between energy and information. bioRxiv .

Stemmler M, Koch C (1999) How voltage-dependent conductances can adapt to maximize the information encoded by neuronal firing rate. Nature neuroscience 2:521–527.

Stemmler M, Sengupta B, Laughlin S, Niven J (2011) Energetically optimal action potentials. Advances in neural information processing systems 24.

Stöber TM, Batulin D, Triesch J, Narayanan R, Jedlicka P (2023) Degeneracy in epilepsy: multiple routes to hyperexcitable brain circuits and their repair. Communications Biology 6:479.

Toni N, Teng EM, Bushong EA, Aimone JB, Zhao C, Consiglio A, van Praag H, Martone ME, Ellisman MH, Gage FH (2007) Synapse formation on neurons born in the adult hippocampus. Nature Neuroscience 10:727–734.

Tuncdemir SN, Lacefield CO, Hen R (2019) Contributions of adult neurogenesis to dentate gyrus network activity and computations. Behavioural Brain Research 374:112112.

Vivar C, Potter MC, Choi J, Lee Jy, Stringer TP, Callaway EM, Gage FH, Suh H, van Praag H (2012) Monosynaptic inputs to new neurons in the dentate gyrus. Nature communications 3:1107.

Whitacre J, Bender A (2010) Degeneracy: A design principle for achieving robustness and evolvability. Journal of Theoretical Biology 263:143–153.

Williams PL, Beer RD (2010) Nonnegative decomposition of multivariate information. arXiv:1004.2515 .

Yang J, Prescott SA (2023) Homeostatic regulation of neuronal function: importance of degeneracy and pleiotropy. Frontiers in Cellular Neuroscience 17:1184563.

Yang K, Sun X, Zhu S (2024) The heterogeneous population of granule cells contributes to pattern separation of the dentate gyrus neural network. Nonlinear Dynamics 112:13465–13481.

Yassa MA, Stark CE (2011) Pattern separation in the hippocampus. Trends in neurosciences 34:515–525.

Yim MY, Hanuschkin A, Wolfart J (2015) Intrinsic rescaling of granule cells restores pattern separation ability of a dentate gyrus network model during epileptic hyperexcitability. Hippocampus 25:297–308.

Young CC, Stegen M, Bernard R, Müller M, Bischofberger J, Veh RW, Haas CA, Wolfart J (2009) Upregulation of inward rectifier k+ (kir2) channels in dentate gyrus granule cells in temporal lobe epilepsy. The Journal of physiology 587:4213–4233.

Yu L, Yu Y (2017) Energy-efficient neural information processing in individual neurons and neuronal networks. Journal of Neuroscience Research 95:2253–2266.

